# STAT5B SH2 variants disrupt mammary enhancers and the stability of genetic programs during pregnancy

**DOI:** 10.1101/2024.05.06.592736

**Authors:** Hye Kyung Lee, Chengyu Liu, Lothar Hennighausen

## Abstract

During pregnancy, mammary tissue undergoes expansion and differentiation, leading to lactation, a process regulated by the hormone prolactin through the JAK2-STAT5 pathway. STAT5 activation is key to successful lactation making the mammary gland an ideal experimental system to investigate the impact of human missense mutations on mammary tissue homeostasis. Here, we investigated the effects of two human variants in the STAT5B SH2 domain, which convert tyrosine 665 to either phenylalanine (Y665F) or histidine (Y665H), both shown to activate STAT5B in cell culture. We ported these mutations into the mouse genome and found distinct and divergent functions. Homozygous *Stat5b^Y665H^* mice failed to form functional mammary tissue, leading to lactation failure, with impaired alveolar development and greatly reduced expression of key differentiation genes. STAT5B^Y665H^ failed to recognize mammary enhancers and impeded STAT5A binding. In contrast, mice carrying the *Stat5b^Y665F^* mutation exhibited abnormal precocious development, accompanied by an early activation of the mammary transcription program and the induction of otherwise silent genetic programs. Physiological adaptation was observed in *Stat5b^Y665H^* mice as continued exposure to pregnancy hormones led to lactation. In summary, our findings highlight that human STAT5B variants can modulate their response to cytokines and thereby impact mammary homeostasis and lactation.

**Graphical abstract:** 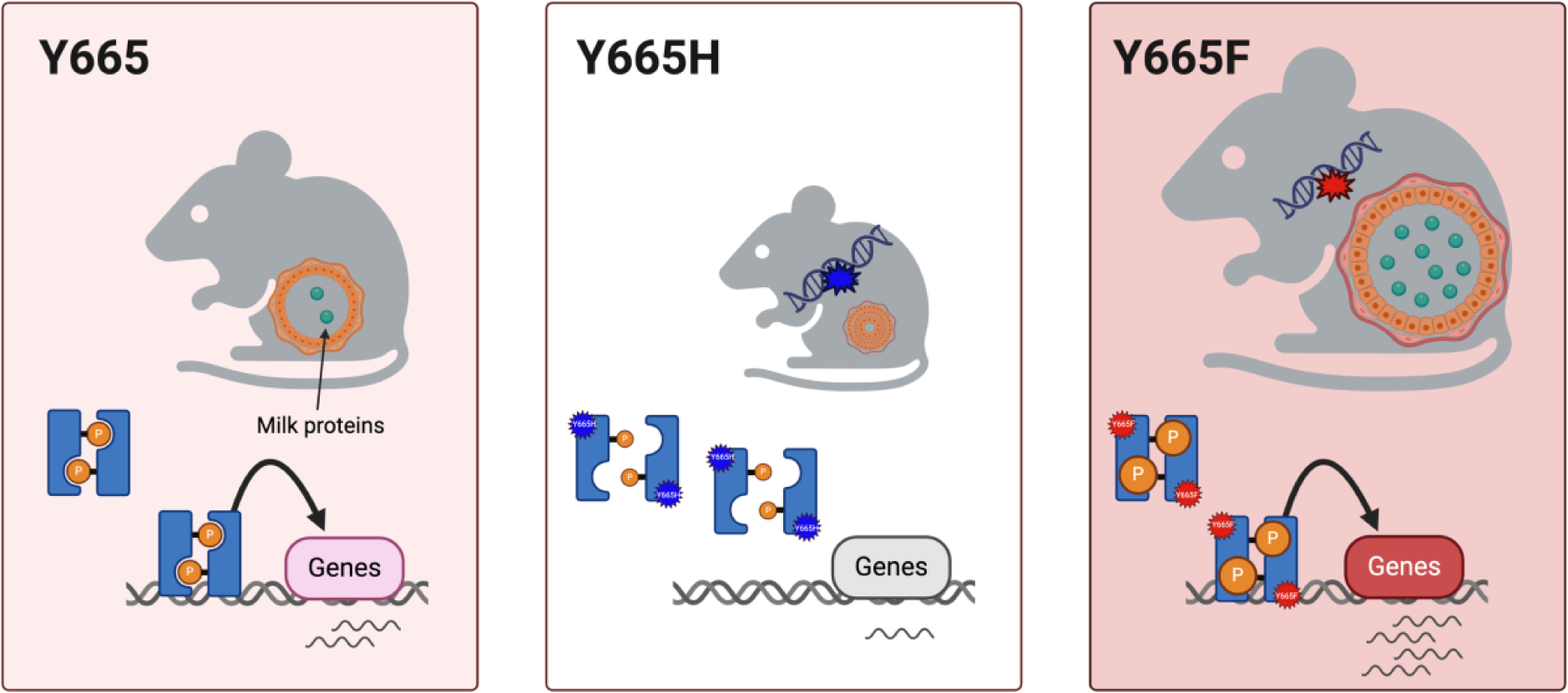

## Introduction

Transcription factors belonging to the Signal Transducers and Activators of Transcription (STAT) family react to various cytokines, initiating both common and cell-specific genetic programs (1). STAT5, the original member of the STAT family, plays a crucial role in cytokine signaling within the hematopoietic system (2), mammary gland (3–5), body growth (6–8), and liver metabolism (8), as evidenced and confirmed through experimental mouse genetics (2,4). Comprising two highly conserved variants, STAT5A (9,10) and STAT5B (10–12), it mediates prolactin signaling during pregnancy, orchestrating the formation and expansion of mammary alveoli to facilitate extensive milk production during lactation, essential for offspring sustenance. Specifically, STAT5A/B bind to precise genomic locations at mammary enhancers and super-enhancers, thereby triggering the induction of milk protein gene expression up to 10,000-fold, during pregnancy and lactation (13–18). While investigations utilizing mouse genetics have shed light on the modulatory role of STAT5A in mammary alveolar differentiation (4), understanding the precise contribution of STAT5B in mammary gland development remains elusive due to the infertility of *Stat5b*-null mice, preventing direct investigation.

Based on available databases, approximately one third of amino acids in the two *Stat5* genes have acquired missense mutations, but genotype–phenotype correlation remains limited. Inactivating mutations in *STAT5B*, associated with Laron syndrome, have been documented in humans (7,19,20), while potential activating mutations have been observed in patients with T cell leukemias (20,21). Since STAT5 responds to cytokine stimuli, the biological consequences of missense mutations might be detected only under specific physiological conditions, such as lactation or a challenged immune system. Although informative, overexpression studies, both in cell lines and mice, may be of limited significance in shedding light on the physiological and pathophysiological functions of missense mutations. A more promising approach involves porting human mutations into the mouse genome, thus permitting investigations on their impact on key target tissues and different physiological states. Development of mammary glands during pregnancy and differentiation during lactation are exquisitely dependent on defined levels of STAT5 (4,5,16), making this organ a perfect test system to investigate the biological impact of naturally occurring STAT5B variants. In this study, we examine the biological consequences of two distinct mutations found in STAT5B, which replace Tyrosine 665 (Y665) with either Phenylalanine (Y665F) or Histidine (Y665H) (20,21). We have ported the Y665F and Y665H missense mutations into the mouse genome and employed lactation, an inherent physiological process in mammalian development, to demonstrate the effects of single amino acid alterations on genetic programs within a hormonally regulated signaling pathway, and how the organism adjusts to these changes. Furthermore, we explore how an activating and inactivating missense mutation in STAT5B influences the function of its twin, STAT5A. This significant example elucidates the fundamental principles of physiological adaptation in response to modifications in transcription factor activity.

## Material and Methods

### Mice

All animals were housed and handled according to the Guide for the Care and Use of Laboratory Animals (8th edition) and all animal experiments were approved by the Animal Care and Use Committee (ACUC) of National Institute of Diabetes and Digestive and Kidney Diseases (NIDDK, MD) and performed under the NIDDK animal protocol K089-LGP-20. CRISPR/Cas9 and base editing targeted mice were generated using C57BL/6N mice (Charles River) by the Transgenic Core of the National Heart, Lung, and Blood Institute (NHLBI). Single-guide RNAs (sgRNA) were obtained Thermo Fisher Scientific’s In Vitro Transcription Service (Supplementary Table 1). Single-strand oligonucleotide donor was obtained from IDT. A plasmid containing the adenine base editor (ABE 7.10 provided by Dr. David Liu’s laboratory) was used to synthesize ABE mRNA using Thermo Fisher’s mMESSAGE mMACHINE T7 In Vitro Transcription Kit. For the Y665H mutant mice, the ABE mRNA (50 ng/μl) and Y665H sgRNA (20 ng/μl) were co-microinjected into the cytoplasm of fertilized eggs collected from superovulated C57BL/6N female mice (Charles River Laboratories). For the Y665F mutant mice, a single-strand oligonucleotide donor contained the desired Y (TAC) to F (TTT) change and a silent C to G change. The silent mutation does not result in amino acid change but can destroy the sgRNA PAM and hence stopping Cas9 from further cutting after the oligo template was successfully knocked in. The Y665F sgRNA was first mixed with Cas9 protein (IDT) to form Cas9 RNP complex, which was co-electroporated with the oligo template into zygotes collected from C57BL/6N mice using a Nepa21 electroporator (Nepa Gene Co) following procedures described by Kenako (22). The microinjected or electroporated zygotes were cultured overnight in M16 medium (Millipore Sigma) at 37°C with 6% CO_2_. Those embryos that reached 2-cells stage of development were implanted into the oviducts pseudopregnant surrogate mothers (Swiss Webster mice from Charles River). All mice born to the foster mothers were genotyped by PCR amplification and Sanger sequencing (Quintara Biosciences) and automate genotyping using a TaqMan-based assay (Transnetyx) with genomic DNA from mouse tails.

Two-months old mice were used in the experiments. Tissues were collected from mice and used immediately or stored at -80°C.

### Whole exome sequencing and data analysis

Genomic DNA was isolated from tail tissue using Wizard Genomic DNA Purification Kit (Promega). Exome sequencing and bioinformatics analyses were performed at Psomage. Target capture for the exome was performed on each sample using SureSelect Mouse All Exon kit (Agilent Technologies). DNA was subjected to SureSelect Target Enrichment System for paired-end DNA library preparation. Whole exome sequencing was performed on a NovaSeq 6000 instrument (Illumina).

Sequencing reads were aligned to the mouse reference (mm10) using BWA 0.7.10. After excluding chimeric reads, the duplicated reads were eliminated using Picard. GATK3.v4 ‘IndelRealigner’ and ‘Table Recalibration’ were used for local realignment and for recalibrating the quality scores, respectively. For SNV/Indel calling in multi-sample analysis, GATK ‘HaplotypeCaller’ was used for comparison with the reference genome. For SNV calling in matched-pair analysis, ‘Selectvariants’ was used to compare the difference between WT and mutant mouse. Annotation for all variants was made using dbSNP142.

### RNA isolation and quantitative real-time PCR (qRT–PCR)

Total RNA was extracted from frozen mammary tissue of wild type and mutant mice using a homogenizer and the PureLink RNA Mini kit according to the manufacturer’s instructions (Thermo Fisher Scientific). Total RNA (1 μg) was reverse transcribed for 50 min at 50°C using 50 μM oligo dT and 2 μl of SuperScript III (Thermo Fisher Scientific) in a 20 μl reaction. Quantitative real-time PCR (qRT-PCR) was performed using TaqMan probes (*Csn1s1*, Mm01160593_m1; *Csn2*, Mm04207885_m1; *Csn1s2a*, Mm00839343_m1; *Csn1s2b*, Mm00839674_m1; *Csn3*, Mm02581554_m1; *Wap*, Mm00839913_m1; *Cish*, Mm01230623_g1; mouse *Gapdh*, Mm99999915_g1, Thermo Fisher scientific) on the CFX384 Real-Time PCR Detection System (Bio-Rad) according to the manufacturer’s instructions. PCR conditions were 95°C for 30s, 95°C for 15s, and 60°C for 30s for 40 cycles. All reactions were done in triplicate and normalized to the housekeeping gene *Gapdh*. Relative differences in PCR results were calculated using the comparative cycle threshold (*C_T_*) method and normalized to *Gapdh* levels.

### Total RNA sequencing (RNA-seq) and data analysis

Ribosomal RNA was removed from 1 μg of total RNAs and cDNA was synthesized using SuperScript III (Invitrogen). Libraries for sequencing were prepared according to the manufacturer’s instructions with TruSeq Stranded Total RNA Library Prep Kit with Ribo-Zero Gold (Illumina, RS-122-2301) and paired-end sequencing was done with a NovaSeq 6000 instrument (Illumina).

Total RNA-seq read quality control was done using Trimmomatic (23) (version 0.36) and STAR RNA-seq (24) (version STAR 2.5.4a) using paired-end mode was used to align the reads (mm10). HTSeq (25) was to retrieve the raw counts and subsequently, R (https://www.R-project.org/), Bioconductor (26) and DESeq2 (27) were used. Additionally, the RUVSeq (28) package was applied to remove confounding factors. The data were pre-filtered keeping only those genes, which have at least ten reads in total. Genes were categorized as significantly differentially expressed with log2 fold change >1 or <-1 and adjusted p-value (pAdj) <0.05 corrected for multiple testing using the Benjamini-Hochberg method were considered significant and then conducted gene enrichment analysis (Metascape, https://metascape.org/gp/index.html#/main/step1). The visualization was done using dplyr (https://CRAN.R-project.org/package=dplyr) and ggplot2 (29).

### Chromatin immunoprecipitation sequencing (ChIP-seq), co-ChIP, and data analysis

The frozen-stored tissues were ground into powder in liquid nitrogen. Chromatin was fixed with formaldehyde (1% final concentration) for 15 min at room temperature, and then quenched with glycine (0.125 M final concentration). Samples were processed as previously described (30). The following antibodies were used for ChIP-seq: STAT5A (Santa Cruz Biotechnology, sc-271542), STAT5B (R&D systems, AF1584; ThermoFisher scientific, 13-5300), H3K27ac (Abcam, ab4729) and RNA polymerase II (Abcam, ab5408). Libraries for next-generation sequencing were prepared and sequenced with the NovaSeq 6000 instrument (Illumina).

For the dimerization of STAT5A/B, chromatin fragments were selectively precipitated by STAT5A antibody, followed by elution facilitated by an IgG elution buffer, pH of 2.8 (Thermo Scientific), and subsequent neutralization employing 1M Tris buffer, pH of 9.5 (Thermo Scientific). The resultant eluted chromatin underwent a secondary precipitation step using STAT5B antibody.

Quality filtering and alignment of the raw reads was done using Trimmomatic(23) (version 0.36) and Bowtie (31) (version 1.2.2), with the parameter ‘-m 1’ to keep only uniquely mapped reads, using the reference genome mm10. Picard tools (Broad Institute. Picard, http://broadinstitute.github.io/picard/. 2016) was used to remove duplicates and subsequently, Homer (32) (version 4.9.1) and deepTools (33) (version 3.1.3) software was applied to generate bedGraph files and normalize coverage, separately. Integrative Genomics Viewer (34) (version 2.3.98) was used for visualization. Coverage plots were generated using Homer (32) software with the bedGraph from deepTools (35) as output. R and the packages dplyr (https://CRAN.R-project.org/package=dplyr) and ggplot2 (27) were used for visualization. Each ChIP-seq experiment was conducted for two replicates and the correlation between the replicates was computed by Spearman correlation using deepTools.

### STAT5B protein structure

The monomeric structure of mouse STAT5B (Uniprot accession P42232) was obtained from the AlphaFold Protein Structure Database (36). Two STAT5B monomer was aligned and then generated a dimer in the crystal structure template of the mouse STAT3 dimer (RCDB ID: 1bg1) (37) using PyMol (38). Additionally, the DNA fragment was coordinated from 1bg1.

### Histology

Fourth inguinal mammary glands tissues from wild-type and *mutant mice* were collected at virgin and days 13 and 18 of pregnancy. Isolated mammary tissues were fixed with 10% neutral formalin solution and dehydrated in 70% EtOH. Samples were processed for paraffin sections and stained with hematoxylin and eosin by standard methods (Histoserv).

### Whole mount and histogram

For whole-mount analysis, an entire mammary gland was flattened on microscopic slides, fixed overnight in Carnoy’s solution (ethanol 6: chloroform 3:acetic acid 1), washed in 70%, 50%, 30%, 10% ethanol and distilled water for 15 min, respectively, and then stained overnight in carmine alum to visualize the ductal trees and alveolar buds. After washing in 70%, 90%, 95%, 100% ethanol, tissues were cleared and stored in xylene. The images were digitized and examined with ImageJ software (version 2.14.0, National Institutes of Health, Bethesda, MD, USA) (39) and histogram was generated using PRISM GraphPad (version 10.1.1).

### Statistical analyses

All samples that were used for qRT-PCR and RNA-seq were randomly selected, and blinding was not applied. For comparison of samples, data were presented as standard deviation in each group and were evaluated with a 1-way or 2-way ANOVA multiple comparisons using PRISM GraphPad. Statistical significance was obtained by comparing the measures from wild-type or control group, and each mutant group. A value of \**p* < 0.05, \*\**p* < 0.001, \*\*\**p* < 0.0001, \*\*\*\**p* < 0.00001 was considered statistically significant. ns, no significant.

## Results

Here we investigate the biological impact of two human STAT5B mutations on the regulation of genetic programs in the mammary gland during pregnancy and lactation. Two distinct SNPs result in the replacement of tyrosine 665 (Y665) within the SH2 domain with either Phenylalanine (Y665F) or Histidine (Y665H) (Figure 1A, B). STAT5B^Y665^ is distinct from STAT5B^Y699^, the key tyrosine required for STAT5B dimerization and nuclear translocation. Y665F replaces the sidechain with a more hydrophobic form of essentially the same type, possibly stabilizing the fold even more, thereby predicting a gain-of-function (GOF). In contrast, Y665H replaces the 6-membered ring with a 5-membered imidazole ring that is positively charged at physiological pH, possibly destabilizing the fold of the monomer, conceivably causing loss-of-function (LOF). Intriguingly, both the Y665F and Y665H mutation result in vastly prolonged activation of STAT5B upon cytokine stimulation (40) suggesting the possibility that they either hyper activate STAT5 target genes or promote aberrant transcription programs normally silent in cells carrying wild type alleles.

**Figure 1.**
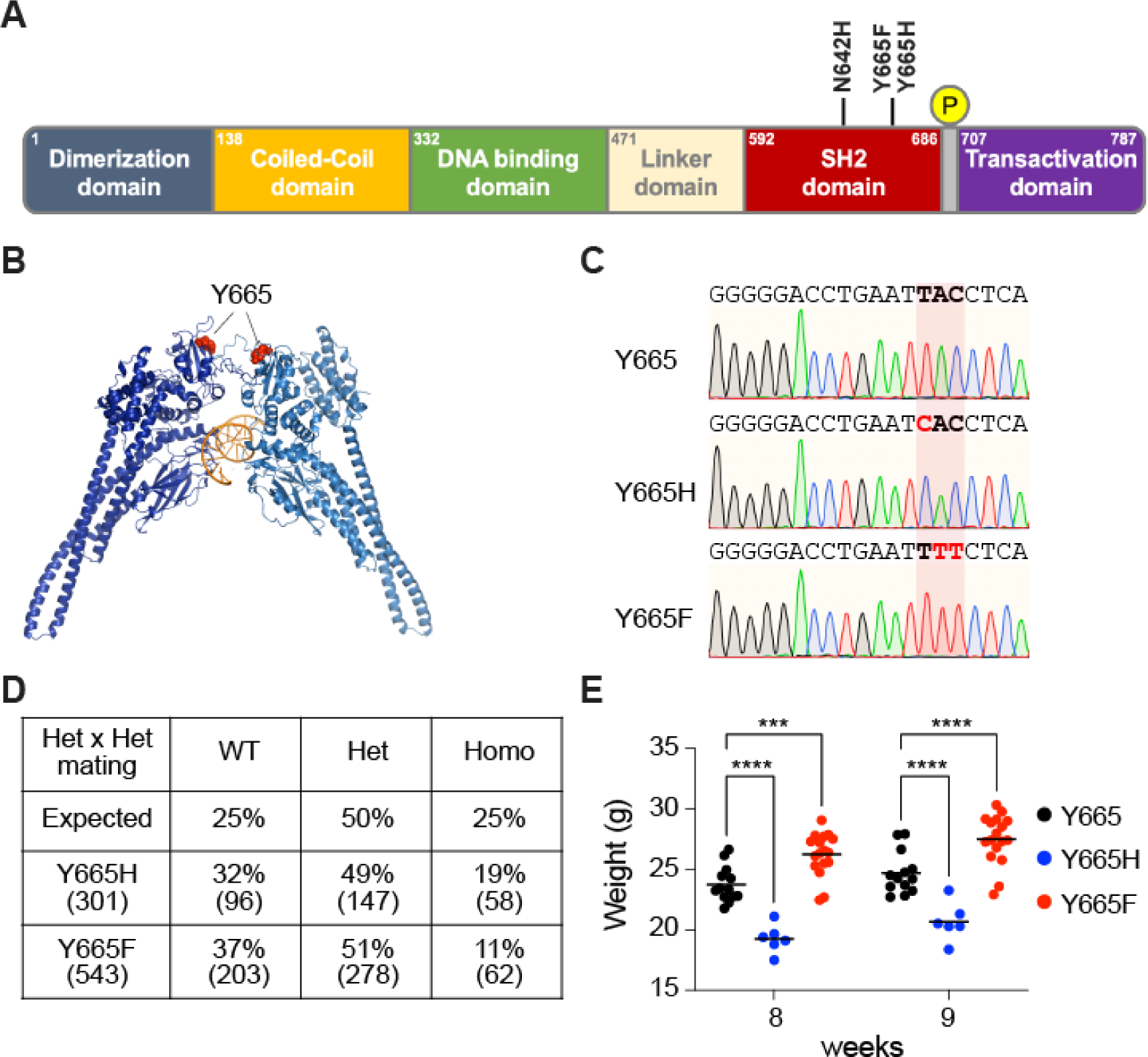
STAT5B mutations at Y665 in the mouse genome. **(A)** Schematic illustration of the STAT5B protein domains. Amino acid locations of the variants are highlighted. **(B)** Three-dimensional model of STAT5B protein structure presenting the location of the Y665 residue mutated in the mouse genome. **(C)** Sanger sequencing chromatograms showing Y665 (wild-type, WT) and the introduction of SNPs resulting in the two missense mutations, *Stat5b^Y665F^* (Y665F) and *Stat5b^Y665H^* (Y665H) mutants. The red shade indicates the altered codons converting Y665 to Y665F and Y665H, respectively. **(D**) Number and percentage of offspring from intercrossed heterozygote parents. **(E)** Dot plot presenting the weight of individual mice in each group from 8 and 9 weeks of age. Results are presented as the means ± SEM of independent biological replicates (Y665, *n* = 13; Y665H, *n* = 6; Y665F, *n* = 17). A two-way ANOVA followed by Tukey’s multiple comparisons test was used to evaluate the statistical significance of differences between groups. \*\**p* < 0.001, \*\*\**p* < 0.00001.

### Introduction of *Stat5b^Y665F^* and *Stat5b^Y665H^* mutations into the mouse genome

To understand the physiological impact of the Y665F and Y665H mutations on hormone-controlled genetic programs, we re-engineered the endogenous *Stat5b* locus using CRISPR/Cas9 and deaminase base editing and generated mice expressing the two *Stat5b* isoforms (Figure 1C). Exome sequencing validated targeting precision and the absence of off-target mutations, including any within the closely related *Stat5a* gene. Lines of mutant mice were established and bred to homozygosity. Only 11% of the pups weaned from *Stat5b^Y665F^* heterozygous crossings were homozygous for the mutation suggesting distinct embryonic or perinatal lethality, while homozygous and heterozygous mice derived from the *Stat5b^Y665H^* heterozygous crossings showed close to expected Mendelian ratios (Figure 1D).

*In vivo*, both mutant mice showed phenotypes clearly demarcating them from wild-type mice. While indistinguishable at the time of weaning, growth curves subsequently diverged in a dimorphic manner at puberty with *Stat5b^Y665F^* mice exhibiting significantly greater body weights and *Stat5b^Y665H^* homozygous mice significantly lower body weights (Figure 1E). These findings establish that the growth hormone signaling cascade, which depends on STAT5B as a key mediator (41), is hyperactivated by the STAT5B^Y665F^ mutation and blunted by the STAT5B^Y665H^ mutation.

### STAT5B^Y665H^ impedes genetic programs in mammary tissue during pregnancy

Although both the STAT5B^Y665F^ and STAT5B^Y665H^ mutation had been identified in human leukemic patients and showed activating features in tissue culture cells, they displayed divergent biological activity when introduced into the mouse genome. While homozygous *Stat5b^Y665H^* females were unable to nurse their litters, *Stat5b^Y665F^* dams successfully raised their pups, suggesting that the Y665H mutation exerts a negative impact on STAT5B function in mammary tissue during pregnancy. However, the Y668H mutation in another isoform, *Stat5a*, did not impact lactation, underscoring the crucial role of STAT5B in mammary differentiation. To understand possible molecular defects imposed by the *Stat5b^Y665H^* mutation, we conducted histological and transcriptional analyses at day 18.5 of pregnancy (p18.5), one day prior to delivery (Figure 2). Histological analyses revealed an undifferentiated mammary alveolar compartment and the absence of extended lumina and milk secretion (Figure 2A). Alveoli were dense without an open lumen and no evidence of any secretory globules. In contrast, mammary alveoli from wild type mice were characterized by open lumina filled with milk fat globules. These findings suggest that the Y665H mutation lacks biological activity to accomplish functional differentiation of mammary alveoli and that the presence of STAT5A fails to compensate for the absence of a functional STAT5B. Notably, mammary tissue from *Stat5b^Y665H^* mice is distinctly different from mice lacking both *Stat5a* and *Stat5b* isoforms (3,5) suggesting that either STAT5A or a residual activity of STAT5B^Y665H^ is sufficient for the establishment of an undifferentiated epithelium. In contrast to *Stat5b^Y665H^*, mammary tissue from *Stat5b^Y665F^* mice had a normal histological appearance (Figure 2A), in line with mutant mice being able to nurse and raise their litters. If anything, *Stat5b^Y665F^* mice displayed a more advanced differentiated alveolar compartment.

**Figure 2.**
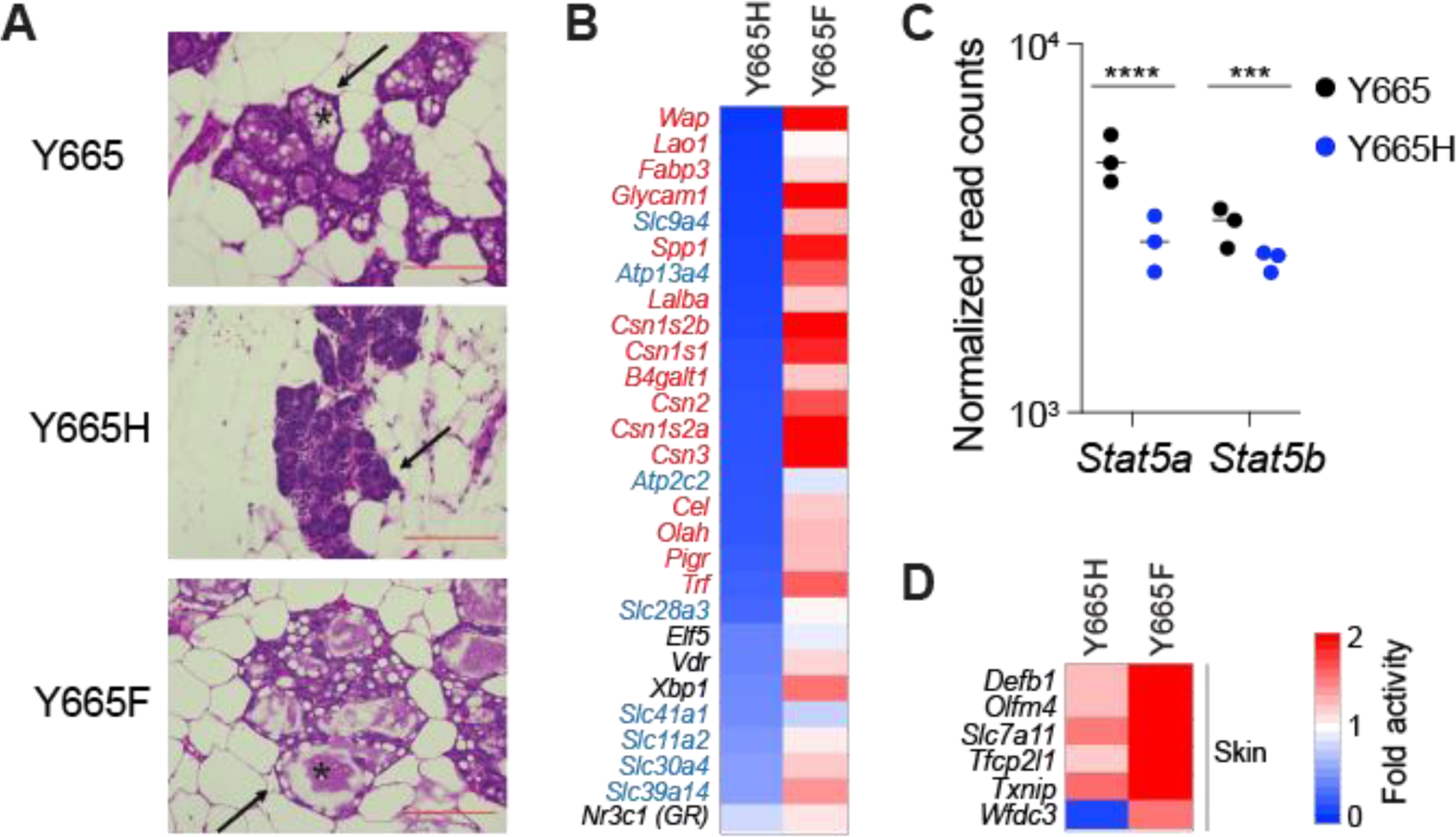
Alveolar development of *Stat5b^Y665^* mutants at day 18.5 of pregnancy. **(A)** Sections of mammary tissue stained with hematoxylin and eosin at day 18.5 with high magnification (x400, Scale bars, 100 μm). The arrow indicates alveoli. **(B and D)** Heatmaps showing fold activity of significantly regulated genes of milk proteins in red, membrane transporters in blue, transcription factors in black (B) and skin-related genes (D) in *Stat5b^Y665H^* (Y665H) and *Stat5b^Y665F^* (Y665F) mutants compared to WT (WT, *n* = 3; Y665H, *n* = 3; Y665F, *n* = 4). **(C)** Dot plots with the normalized read counts to mRNA levels of *Stat5* genes controlled by autoregulatory enhancer. Results are shown as the means ± SEM of independent biological replicates (*n* = 3). The Benjamini-Hochberg adjusted *p*-value was used for significance. \*\*\**p* < 0.0001, \*\*\*\**p* < 0.0001.

Genes encoding milk proteins and other components controlling milk production and secretion during lactation, such as membrane transporters and enzymes, are activated up to several-thousand-fold during pregnancy, a process largely controlled by the combined action of JAK2 (18) and STAT5A/B (16). To determine to what extent expression of these genes was impacted by the two *Stat5b* variants, Y665H and Y665F, we conducted RNA-seq from mammary tissue collected at day 18.5 of pregnancy, just prior to parturition. Expression of approximately 660 genes was reduced by at least 50% in *Stat5b^Y665H^* mice (Supplementary Table 2). To gauge the direct impact of STAT5B^Y665H^ on mammary epithelial differentiation, we focused on three functional gene classes, milk protein genes, membrane transporters contributing to milk secretion and regulatory proteins (receptors, transcription factors, enzymes). Expression of milk protein genes, that comprise more than 95% of all mRNAs, was greatly reduced, ranging from a more than a 90% decline for genes, such as *Wap* and *Glycam1*, to a more modest 50% for membrane transporters (Figure 2B). STAT5A/B not only controls the expression of milk protein, but also activates transcription factor genes, such as *Elf5*, *Vdr*, *Xbp1* and *Nr3c1* that are known to participate in the activation of mammary genetic programs. Expression of these TFs was reduced by 50-60% (Figure 2B). Autoregulation of the *Stat5a/b* locus is a notable feature that facilitates its critical role in the activation of mammary genetic programs (30) and reduced *Stat5a* expression was observed in *Stat5b^Y665H^* mice (Figure 2C). Lastly, expression of members of the solute carrier (SLC) protein family is greatly reduced, supporting the absence of open mammary alveolar lumina and milk in *Stat5b^Y665H^* mice (Figure 2B). These findings reinforce that STAT5B^Y665H^ has an impaired transcriptional activity that cannot be fully compensated for by the presence of STAT5A. Moreover, the mammary-centric transcription factor NFIB (42) might support the observed remaining low expression of mammary genes.

### STAT5B^Y665F^ results in exacerbated mammary differentiation during pregnancy

In contrast to *Stat5b^Y665H^*, *Stat5b^Y665F^* mice exhibited exacerbated mammary differentiation at day 18.5 of pregnancy as evidenced by histological appearance (Figure 2A). This was further supported by an elevated expression of some milk protein genes, such as *Csn2* (Figure 2B; Supplementary Table 3). However, this increase was less than 2-fold, in sharp contrast to the 95% reduction seen for some genes in *Stat5b^Y665H^* mice. Unexpectedly, STAT5B^Y665F^ activated a set of approximately 140 genes that are normally not under the control by pregnancy hormones through JAK-STAT. Among those, *defensin beta 1* (*Defb1*) is activated more than 17-fold in *Stat5b^Y665F^* mice (Figure 2D). These findings suggest that the STAT5B^Y665F^ mutation has a modest positive impact on *bona fide* mammary-specific genes but aberrantly activates gene classes that are not under pregnancy control, including genes key for skin homeostasis.

### STAT5B^Y665F^ and STAT5B^Y665H^ differentially impact mammary enhancers

Gene classes induced during pregnancy are most often activated by STAT5-based enhancers and super-enhancers (13–16), begging the question if STAT5B^Y665F^ and STAT5B^Y665H^ impact the enhancer landscape and thereby expression of STAT5 target genes. Specifically, we asked if the STAT5B^Y665H^ mutant displays compromised binding to GAS motif, thereby failing to activate mammary enhancers. Conversely, we asked if increased target gene expression in *Stat5b^Y665F^* mice was the result of enhanced binding to enhancers. Towards this end, we conducted ChIP-seq experiments and investigated the capacity of STAT5B^Y665H^ and STAT5B^Y665F^ to recognize GAS motifs (TTCnnnGAA) within enhancers linked to mammary-specific and more widely expressed STAT5 target genes (Figure 3). While STAT5B binding to mammary-specific enhancers, such as those driving the *Casein* locus (15) or the *Wap* gene (*13*) was detected in tissue collected from wild type mice at day 18.5 of pregnancy, little or no binding was observed in tissue collected from *Stat5b^Y665H^* mice. The absence of STAT5B binding paralleled reduced H3K27ac coverage and Pol II loading (Figure 3A), supporting their reduced expression levels. As outlined above, the remaining expression of milk protein genes observed in *Stat5b^Y665H^* mice might be the result of other transcription factors, such as NFIB, known to bind to mammary enhancers (13,15,42), a concept we tested using ChIP-seq experiments. Notably, NFIB binding to the *Casein*, *Wap*, *Slc9a4* and *Xdh* enhancers was not significantly reduced in *Stat5b^Y665H^* mice (Figure 3A), establishing NFIB as a transcription factor that can function in the presence of a transcriptionally impaired STAT5B.

**Figure 3.**
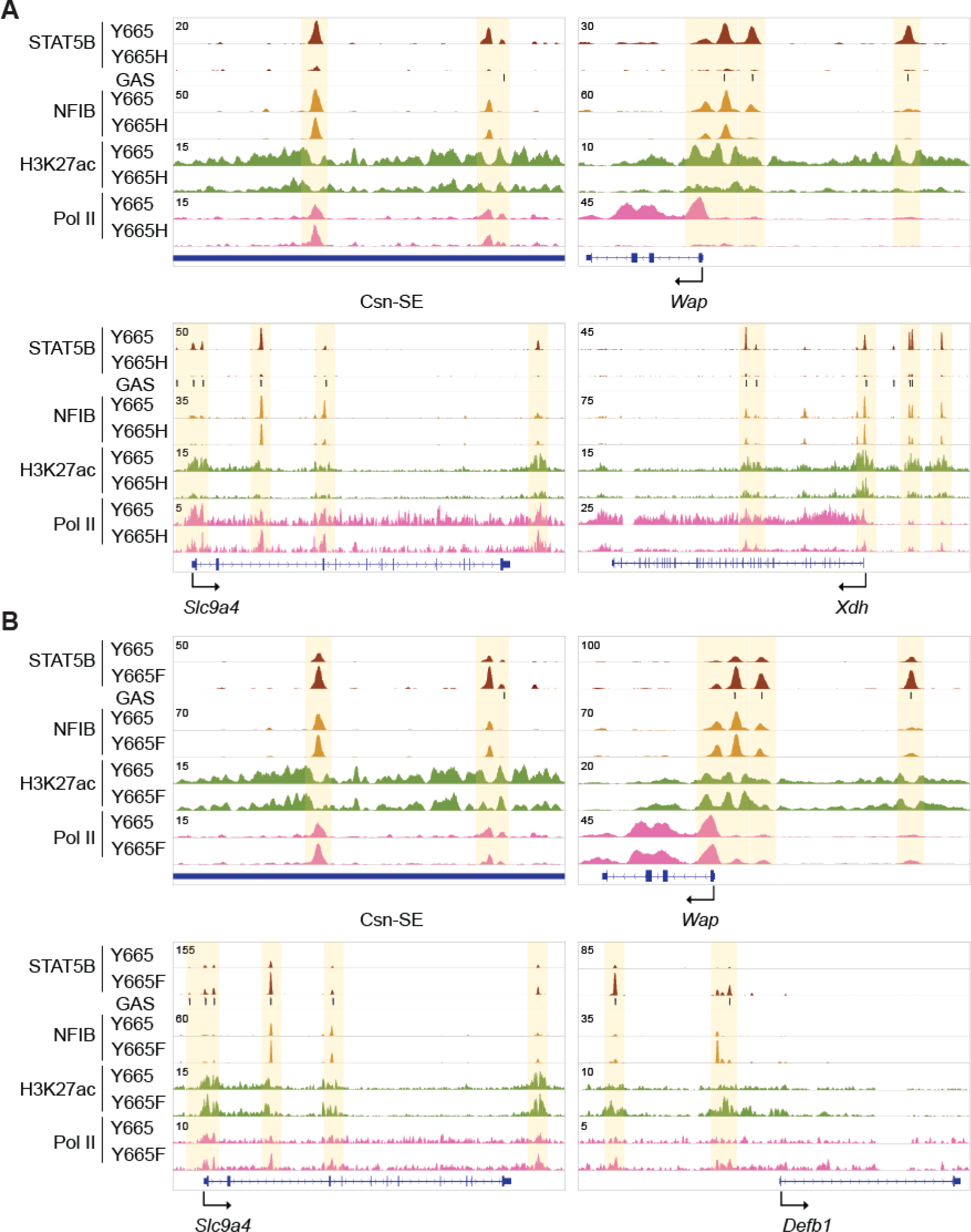
Genomic features of the genes regulated by *Stat5b^Y665^* mutations. **(A)** Binding of STAT5B and NFIB, H3K27ac, and Pol II at gene loci of milk protein genes controlled by mammary-specific super-enhancers, transporter genes, and skin genes in mammary tissue of wild type (Y665) and *Stat5b^Y665H^* (Y665H) mice. **(B)** STAT5B and NFIB binding, H3K27ac and Pol II loading at gene loci of milk protein genes controlled by mammary-specific super-enhancers, transporter gene and skin gene in mammary tissue of Y665 and *Stat5b^Y665F^* (Y665F) mice.

Next, we examined whether the gene activation observed in *Stat5b^Y665F^* mice could be explained by an enhanced STAT5B binding to mammary enhancers and super-enhancers. Increased STAT5B binding to known mammary enhancers was observed in *Stat5b^Y665F^* mice (Figure 3B). Importantly, strong STAT5B binding was observed at candidate enhancers of the *de novo* induced genes, such as *Defb1* (Figure 3B) whose expression is induced 17-fold. Concordantly, NFIB binding was enhanced suggesting either an indirect recruitment through STAT5B or binding to NFIB motifs in a more optimal accessible chromatin.

### Temporal developmental shift of mammary differentiation in *Stat5b^Y665F^* mice

Genes encoding milk proteins and other regulatory proteins display distinct temporal activation patterns in mammary tissue with a sharp increase of transcription frequently observed in the latter part of pregnancy (16,17). Since STAT5B^Y665F^ is a *bona fide* activating mutation, we asked if it would lead to a temporal shift of the differentiation program during pregnancy. For this, we investigated the histological appearance and transcription programs of mammary tissue at day 13.5 of pregnancy (p13.5), a stage when key mammary genes have not been activated due to suboptimal STAT5 signaling. Histologically, STAT5B^Y665F^ tissue displayed an advanced degree of development and differentiation as indicated by an accumulation of lipid droplets and open lumina (Figure 4A). Transcriptome analysis revealed a sharp increase of milk protein gene transcripts, such as *Spp1* and *Wap*, the mammary-centric transcription factor *Elf5* and membrane transporters required for secretion of milk (Figure 4B; Supplementary Table 4). Expression of some of these genes approached levels seen at day 18.5 of pregnancy (p18.5) in wild type (Figure 4C), confirming precocious differentiation. ChIP-seq experiments confirmed increased binding of STAT5B^Y665F^ to key mammary enhancers (Figure 4D).

**Figure 4.**
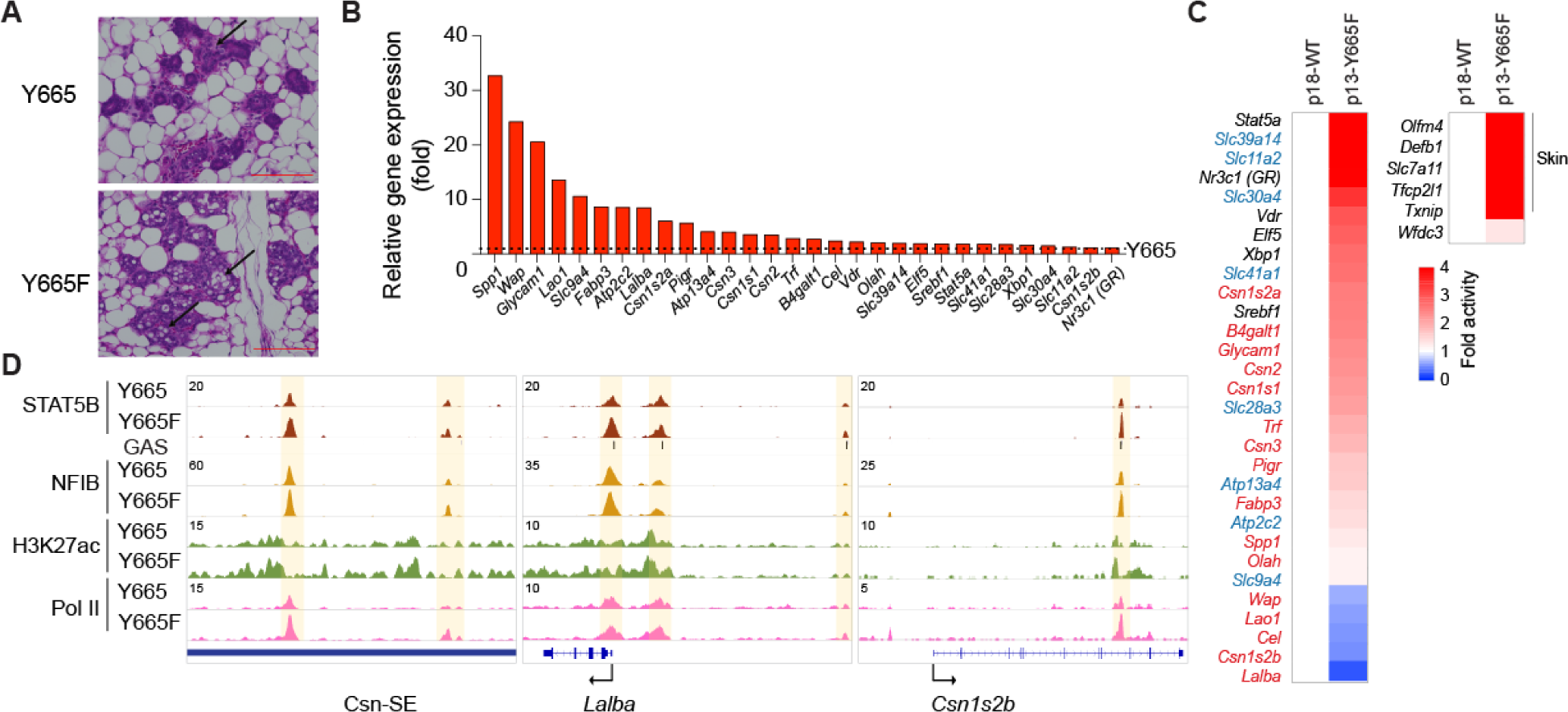
Alveolar development of *Stat5b^Y665F^* mutants at mid-pregnancy. **(A)** Sections of mammary tissue stained with hematoxylin and eosin at day 13.5 with high magnification (x400, Scale bars, 100 μm). **(B)** Bar graphs showing fold activity of genes of milk proteins, transporters and transcription factors in *Stat5b^Y665F^* (Y665F) mice compared to wild type (Y665) (WT, *n* = 3; Y665F, *n* = 4). **(C)** Heatmaps presenting fold activity of milk protein genes in red, transporter genes in blue, transcription factors in black (left) and skin genes (right) in *Stat5b^Y665F^* (Y665F) mice at day 13.5 of pregnancy (presented as p13) compared to WT at day 18.5 of pregnancy (presented as p18) (WT at p18, *n* = 3; Y665F at p13, *n* = 4). **(D)** Binding of STAT5B and NFIB, H3K27ac, and Pol II at gene loci of milk protein genes controlled by mammary-specific super-enhancers in mammary tissue of Y665 and Y665F mice at day 13.5 of pregnancy.

Having observed that mammary alveolar differentiation is shifted to mid pregnancy in *Stat5b^Y665F^* mice, we asked whether the activated STAT5B also induces mammary-specific genes in non-parous mice, i.e. in the absence of continuous prolactin stimuli (Figure 5). First, we investigated the density of mammary structures, both ducts and alveoli, using mammary whole mounts (Figure 5A-B). Density was increased in *Stat5b^Y665F^* mice and decreased in *Stat5b^Y665H^* as compared to wild type mice, suggesting that the two mutants impacted mammary development already in the absence of a pregnancy-induced hormonal milieu. Importantly, expression of key milk protein genes increased more than 30-fold in *Stat5b^Y665F^* mice, while their expression decreased in *Stat5b^Y665H^* mice. Expression of *Cish*, a STAT5-controlled genes expressed across many cell types, was unaltered (Figure 5C).

**Figure 5.**
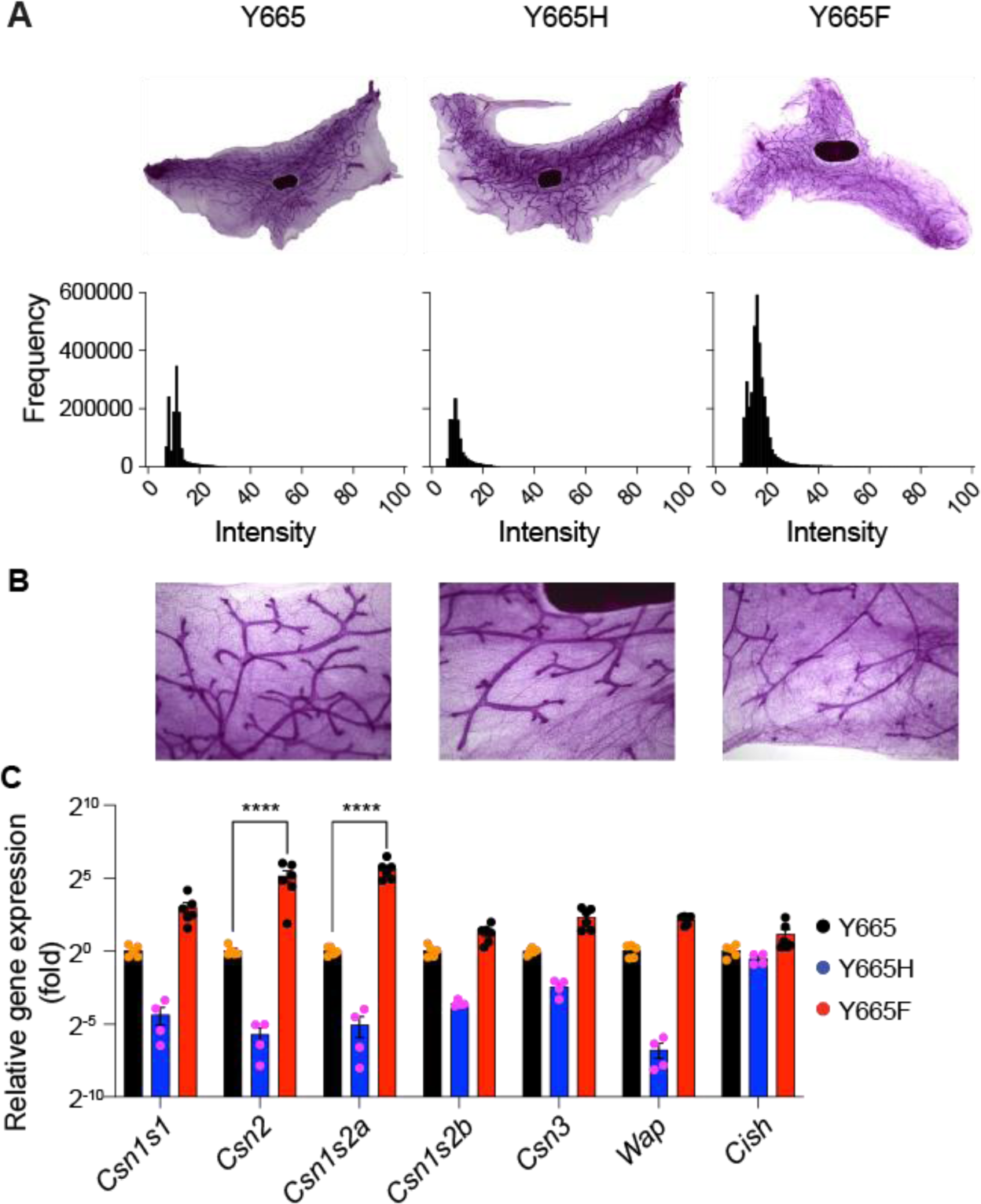
Mammary gland structures in virgin tissue of mutants. **(A)** Image of carmine alum-stained whole mount in virgin mammary gland of WT and mutant mice and histogram processed by Gaussian denoising. **(B)** Zoom-in image of each whole mount image. **(C)** Expression of representative milk protein genes (*Casein* and *Wap* genes) was measured in mammary tissue of Y665 (WT) and mutant mice collected at virgin by qRT- PCR (WT, *n* = 5; Y665H, *n* = 4; Y665F, *n* = 6). Results are shown as the means ± SEM of independent biological replicates. *p*-values are from 2-way ANOVA followed by Dunnett’s multiple comparisons test between WT and mutants. \*\*\*\**p* < 0.00001.

### Physiological adaptation of *Stat5b^Y665H^* mice after subsequent pregnancies

Although prolactin exposure during the 18.5 days of the first pregnancy was not sufficient to build functional mammary glands in *Stat5b^Y665H^* mice, it was unclear whether continuous hormonal stimulation would overcome the developmental block. To test this, we mated *Stat5b^Y665H^* homozygous mice immediately after their first pregnancy, thereby enabling a continued prolactin stimulus on mammary epithelium. STAT5B^Y665H^ dams were able to support litters after their second pregnancy, although the pups grew slower (Figure 6A), suggesting reduced milk production. To gauge the differentiation status of mammary tissue, and thereby indirectly milk protein production, we measured mRNA levels of distinct milk protein mRNAs at the end of the first pregnancy (p18.5) and at day 10 of lactation (L10) after the second pregnancy (Figure 6B). While the expression of key milk protein genes was reduced by up to 90% at the end of the first pregnancy of STAT5B^Y665H^ dams as compared to control mice carrying wild type STAT5B, expression levels increased several-fold after the second pregnancy. Expression of these milk protein was at least 50% of that seen in control mice (Figure 6C) supporting the ability of the STAT5B^Y665H^ dams to raise their litters. As anticipated from data at earlier stages, *Stat5b^Y665F^* mutant exhibited heightened levels of milk protein genes at L10.

**Figure 6.**
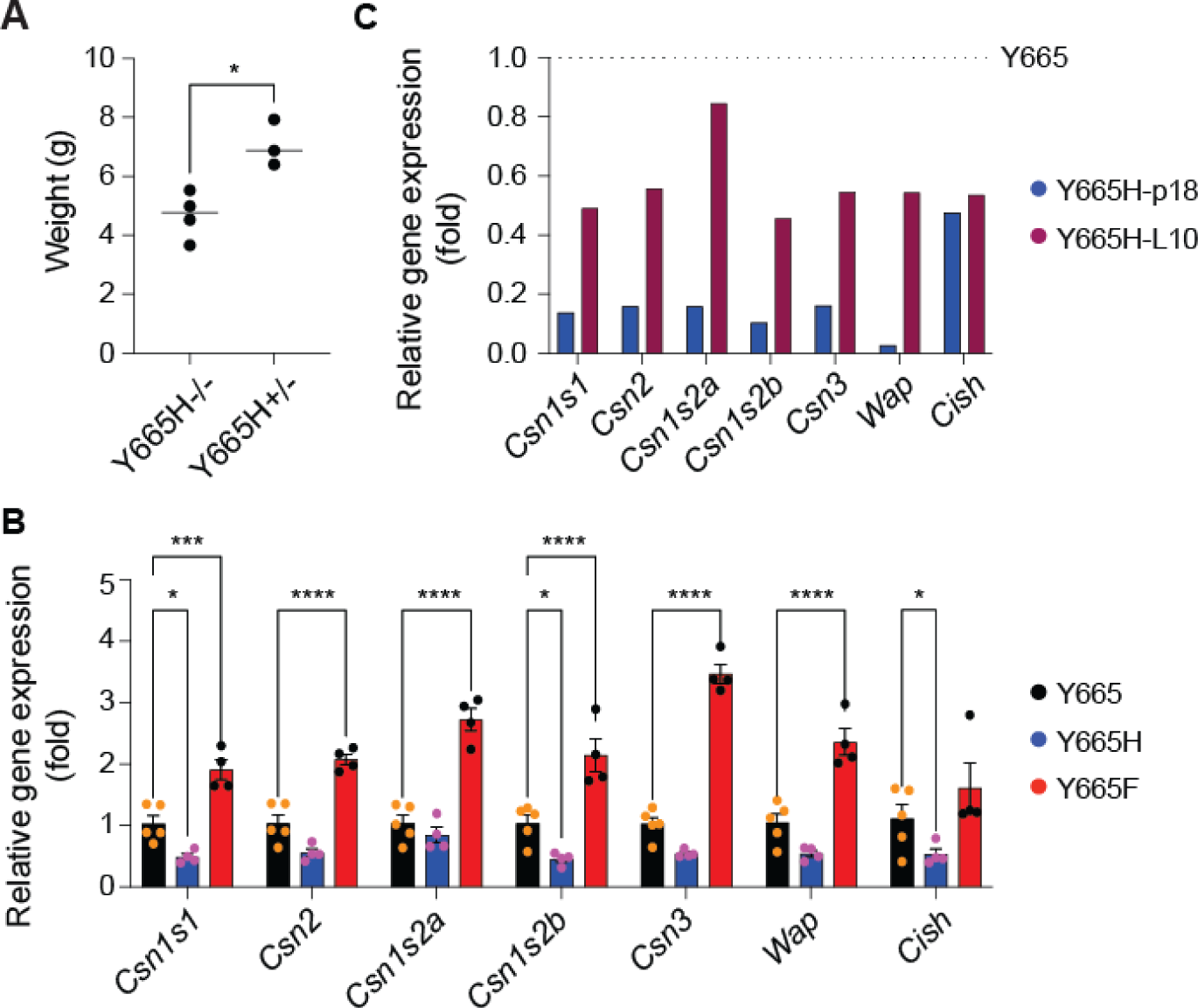
Partial activation of mammary function and lactation in STAT5B^Y665H^ mice after the second pregnancy. **(A)** Body weight of pups at day ten of lactation (L10) after the second pregnancy of Y665H^-/-^ and after the first pregnancy of Y665H^+/-^. The weight of pups from 3-4 females was measured and normalized by the number of pups. A *t*-test was utilized to evaluate the statistical significance between pups from homozygous females at second pregnancy and heterozygous females at first pregnancy. \**p* < 0.05. **(B)** Expression of representative milk protein genes (*Casein* and *Wap* genes) was measured in mammary tissue of Y665 (WT) and Y665F mice collected at day 10 of lactation (L10), and of Y665H mice at L10 after the second pregnancy by qRT-PCR (Y665, *n* = 5; Y665H, *n* = 4; Y665F, *n* = 4). Results are shown as the means ± SEM of independent biological replicates. *p*-values are from 2-way ANOVA followed by Dunnett’s multiple comparisons test between WT and mutants. \**p* < 0.05, \*\*\**p* < 0.0001, \*\*\*\**p* < 0.00001. **(C)** The relative reduction rate of representative milk protein genes and the common STAT target gene (*Cish*) between WT and Y665H mice at p18 and L10 was presented by a bar graph.

### STAT5A/B dimerization is impaired in *Stat5b^Y665H^* mice

STAT proteins are known to form homodimers and heterodimers (43), permitting diversity of transcriptional responses. We had originally hypothesized that STAT5A would compensate, at least in part, for the absence of a functional STAT5B. However, the greatly impaired expression of milk protein genes in *Stat5b^Y665H^* mice suggested the inability of STAT5A to compensate for the defective STAT5B. The possibility arose that STAT5A cannot activate mammary target genes by itself or, alternatively, STAT5A binding to its target sites in enhancers is dependent on a functional STAT5B. To address these options, we first determined if STAT5A and STAT5B form heterodimers in mammary tissue. For this we conducted a two-step ChIP-seq experiment. In the first step, chromatin was precipitated using STAT5A-specific antibodies, followed by elution. Secondly, the eluted chromatin, precipitated by the STAT5A protein, was then subjected to precipitation again using STAT5B-specific antibodies (Figure 7A). This experiment demonstrated that STAT5A/B heterodimers occupy mammary-specific and common enhancers (Figure 7B).

**Figure 7.**
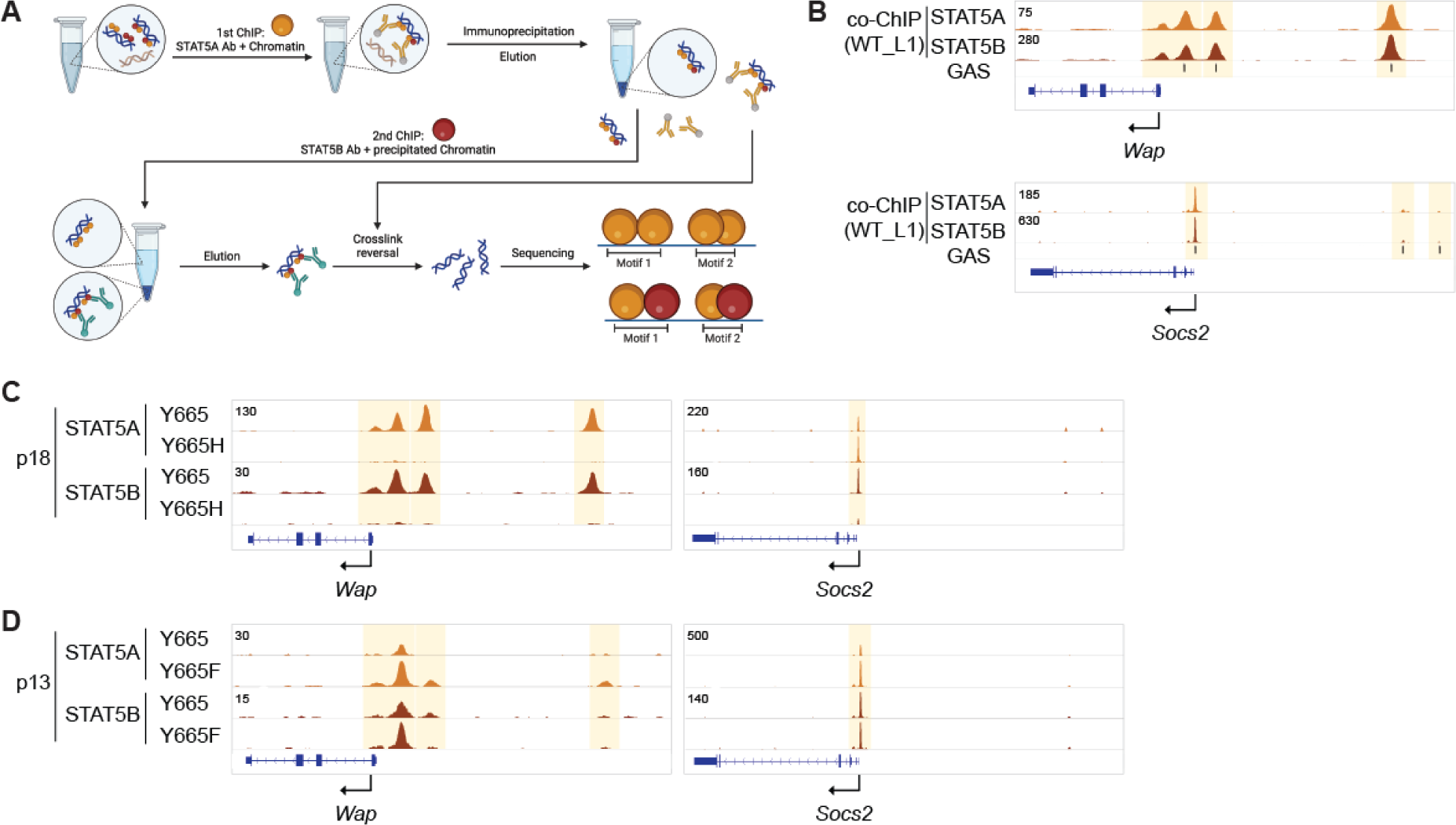
Dimerization of STAT5A and STAT5B protein. **(A)** Strategy of co-ChIP-seq. STAT5A bound chromatin from mammary tissue of WT mice at day 1 of lactation (L1) was precipitated using STAT5A antibody and eluted from Ig beads. Secondly, STAT5B bound chromatin was precipitated using STAT5B and eluted from the beads. STAT5A motifs from 1st ChIP and STAT5B motifs from 2nd ChIP were identified by sequencing. **(B)** Co-ChIP profile at the representative milk protein *Wap* gene and common STAT target *Socs2* gene. **(C)** STAT5A and STAT5B binding at gene loci of milk protein *Wap* gene and common STAT target *Socs2* gene in the mammary gland of Y665 (WT) and Y665H mice at day 18.5 of pregnancy (p18). **(D)** STAT5A and STAT5B binding in the mammary gland of Y665 (WT) and Y665F mice at day 13.5 of pregnancy (p13).

Next, we investigated if STAT5A binding was impacted by the STAT5B^Y665H^ mutant and conducted ChIP-seq experiments from mammary tissue harvested at day 18.5 of pregnancy (**Figure 7C**). Notably, in addition to the failure of STAT5B^Y665H^ to bind to mammary enhancers, no STAT5A binding was detected in the *Wap* enhancers, suggesting that STAT5A/B heterodimers are the major complex recognizing these enhancers. In contrast, STAT5A binding to common STAT5-controlled enhancers in *Socs2* was not visibly affected (Figure 7C). Based on these results it could be predicted that enhanced STAT5B^Y665F^ activity would recruit additional STAT5A to enhancers. To test this, we conducted STAT5A ChIP-seq in STAT5B^Y665F^ tissue at day 13.5 of pregnancy (Figure 7D). Elevated STAT5A occupancy was detected at mammary enhancers in *Stat5b^Y665F^* mice, as indicated at the *Wap* gene, as compared to control tissue.

## Discussion

The insertion of human mutations into the mouse genome permitted a robust comparative investigation and uncovered unexpectedly intricate properties of two human STAT5B variants that had previously been identified in T cell leukemias. It uncovered the essential function of tyrosine 665 within the STAT5B SH2 domain in mammary gland homeostasis and lactation, and unique functions of two missense mutations replacing this tyrosine. Our study exposed that the two mutations elicited distinctly opposite effects on transcription enhancers and super-enhancers resulting in altered genetic programs and mammary development and lactation. While the phenylalanine residue enhanced STAT5B activity, the histidine residue resulted in an inactive STAT5B. Analysis of mammary tissue showed that the behavior of the inactivating mutation varied markedly between the first and second pregnancy, emphasizing the contribution of continued stimulation by pregnancy hormone in overcoming partially inactivating mutations and attaining functional mammary development and lactation.

The two mutations represent the two classes of activation and inactivation. Unexpectedly, the Y665H mutant strongly echoed expectations of an inactive STAT5B, while the Y665F mutation displayed intensified activity. Both mutations had initially been identified in human leukemic patients (20,21), both demonstrating activation phenotypes in tissue culture cell experiments (40). The apparently contradictory results on the Y665H mutation may be secondary to the differences in experimental approaches, here the use of gene editing to insert of the mutation into the endogenous *Stat5b* locus while the tissue culture cell experiments were based on retroviral mediated overexpression. The functional impact of the two mutations *in vivo* highlight the role of STAT5B in mediating sexually dimorphic growth, the activating mutation accelerating and the inactivating one retarding growth in males (41).

The full *in vivo* significance was revealed by examining the interplay of organ growth and transcriptional activation occurring with maximal physiological stimulation of STAT5 signaling through prolactin signaling. Y665H blunted mammary development and STAT5 transcriptional activation while Y665F mediated precocious mammary development in pregnancy and augmented gene expression of STAT5 target genes. Mechanistically, diminished STAT5B function resulted in failure to establish functional enhancer complexes, thereby limiting transcriptional initiation of milk protein and other genes required for milk production such as transporters. Importantly, while a single round of pregnancy failed to activate the Y665H mutant, repeated pregnancy enabled partial activation with sufficient milk protein gene expression to sustain lactation and pup survival. This illustrates the power of functional adaptation in, at least partially, overriding inactivating STAT5 mutations (44).

In the Y665F mice variant alveolar differentiation was temporally shifted towards mid as compared to late pregnancy as is normally observed. Significantly, even non-parous mice showed differences in mammary gland growth with the Y665F variant, likely secondary to exaggerated growth during repetitive estrous cycles mediated by prolactin. Functionally, the precocious growth likely reflects precocious activation of key mammary enhancers and super-enhancers at early stages of pregnancy as well as during specific phases of the estrous cycle in virgin mice. It would appear that hyperactive variants, such as Y665F can access native chromatin that is otherwise inactive due to its methylation and acetylation status (45). Transcriptional analyses showed that Y665F not only preferentially hyperactivated normally activate genetic programs, but it also induced genes that are silent during normal development. This finding demonstrates that activating STAT5B variants can induce aberrant genetic programs with implications for mammary cancer and immune homeostasis (46). Although no naturally occurring human STAT5A activating mutations have been identified, experimentally designed activating mutations such as *Stat5a^S711F^* (47,48) is highly oncogenic when introduced into transgenic mice (49–51) and induces pregnancy-independent mammary development following introduction by viral vectors (52).

STAT5A, originally called mammary gland factor (MGF) (9), is critical for normal mammary differentiation during pregnancy and lactation (4). Specific study of the contribution of STAT5B to this process has been hampered by the infertility of STAT5B knockout mice (41). An important contribution of this study is to define STAT5B as the more critical STAT5 gene in driving pregnancy-mediated mammary development, highlighting its important function as a heterodimer with STAT5A. A preferential heterodimerization of STAT5A/STAT5B may represent a tissue specific or developmentally mediated function as under *in vitro* condition in T cells this is not observed (53,54).

STAT protein SH2 domains support their JAK2-mediated phosphorylation, dimerization, nuclear translocation and binding to regulatory elements in promoters and enhancers. To date, missense mutations have been identified at approximately 30 amino acid positions within the STAT5B SH2, falling into three categories: gain-of-function (GOF), loss of function (LOF) and dominant negative. STAT5B SH2 mutations associated with the growth hormone insensitivity (GHI) syndrome include A630P, K632N, F646S and V669F result in postnatal growth failure and, in some cases, immune deficiency (7,55,56). They are considered to disrupt the anti-parallel β-sheets and thereby the pocket for binding phosphate groups on respective receptors, resulting in abrogated STAT5 activation. The Y665H variant studied here would also be predicted to result in disruption of the anti-parallel β-sheets and abrogation of STAT5B function, observed experimentally here.

Expanding transcriptional programs through activating mutations in transcription factors is a well-recognized theme traceable to different domain structures. GOF mutations in STAT3 have been identified in DNA binding, coiled-coil, and SH2 domains. Deregulated transcriptional programs have been identified in immune cells of mice carrying GOF STAT3 human variants (57–60). Because we saw similar magnitudes of induction in transcriptional programs with Y665F, we conjecture that viable activating mutations may have similar capacities across different STATs and cell types. A general theme might be that GOF mutations outside the DNA binding domain preferentially hyperactivate existing programs while mutations in DNA binding domains could yield the emergence of novel and unanticipated programs. The C99R mutation in the DNA binding domain of the Interferon Regulatory Factor 4 (IRF4), a key TF in immune cells, redirects binding to different motifs and therefore fundamentally alters transcriptional output (61).

Women who experience suboptimal milk production with a first pregnancy can find that lactation improves following the second pregnancy (62). Although the genetic basis for this phenomenon has not been widely investigated, mutations in the prolactin gene have been identified in women with puerperal alactogenesis (63). These inactivating mutations would inevitably result in a suboptimal activation of STAT5 and subsequent difficulties in maintaining lactation. Additional Individuals with prolactin gene mutations can be identified in general population databases such as AllofUs, providing the opportunity to investigate the role of genetic variations in lactational diversity. In mice, an activating mutation in the viral sensor 2’-5’- oligoadenylate synthetase (*Oas*) 2 gene leads to lactational failure (64) in association with induction of immune pathway genes in mammary alveolar epithelium. Since *Oas2* is a STAT5 target (65,66), it can be hypothesized that recruitment of other genes under STAT5 control in mammary alveolar cells could impact lactational performance. Mutational studies in mice have shown that the deletion of a key mammary super-enhancer (15) and the enhancer driving the *Csn3* gene (15) resulted in reduced lactational performance. Similarly, several metabolic genes under STAT5 control are critical for successful lactation (67). It is possible more information on the genetic of lactation could be revealed by further study of individuals with inactivating and dominant negative STAT5B mutations that disrupt the growth hormone axis and immune homeostasis (68).

In summary, this is the first report to demonstrate that human variants in the JAK-STAT signaling pathway significantly disrupt mammary physiology and lactation. The increasing number of publicly available human genetic databases with associated physiological and medical information such as AllofUs provides an opportunity to investigate the role of genetic variation in lactational diversity in women.

## Limitations of the study

At this point, the reason for the distorted Mendelian ratio, i.e. the paucity of Y665F homozygous mice derived from mating heterozygous females with heterozygous males is not known. Also, the molecular mechanisms through which the inactivating Y665H mutation gains limited function after the second pregnancy remains unknown.

## Data availability

All data will be uploaded to Gene Expression Omnibus (GEO).

## Author contributions

H.K.L. and L.H. jointly designed the study and analyzed data. C.L. generated mutant mice. H.K.L. established mutant mouse lines, performed experiments and data analysis. H.K.L. and L.H. wrote the manuscript and all authors approved the final version.

## Acknowledgements

We thank the NIDDK genomics core and the NHLBI sequencing core for NGS, Sung-Gwon Lee for advice in bioinformatic analysis and Priscilla A. Furth for discussion and advice. This work utilized the computational resources of the NIH HPC Biowulf cluster (http://hpc.nih.gov).

## Funding

H.K.L. and L.H. were supported by the Intramural Research Programs (IRPs) of National Institute of Diabetes and Digestive and Kidney Diseases (NIDDK) and C.L. was supported by Intramural Research Programs (IRPs) of National Heart, Lung, and Blood Institute (NHLBI).

## Conflict of interest

The authors have no competing interests.

## Notes

### Competing Interest Statement

The authors have declared no competing interest.

